# Identification of Human Pluripotent Stem Cell Derived Astrocytic Progenitors that Correlate with Glioblastoma Subtypes

**DOI:** 10.1101/2025.08.26.672421

**Authors:** Stephanie Van Gulden, Anne Kathryn Linden, John A. Kessler, Chian-Yu Peng

## Abstract

Recent studies have identified stem/progenitor cells in Glioblastoma multiforme (GBM) tumors that recapitulate developmental glial lineages in tumor progression, but whether non-malignant developmental glial progenitors share molecular characteristics with GBM tumor subtypes has not been extensively investigated. Here we present an approach that uses human pluripotent stem cells (hPSCs)-derived neural progenitors to study the developmental diversity of cells in the astrocytic lineage and how it correlates with GBM subtypes. Using a combination of single cell RNA sequencing of gliogenic stage neural progenitors and bulk RNA sequencing of cells that express combinations of GBM-associated cell surface markers, we identified two astrocytic progenitor populations that share gene expression profiles with the mesenchymal and proneural GBM subtypes respectively. Differential gene expression and pathway analyses of mesenchymal and proneural-associated progenitor clusters revealed enrichment in TGF-β signaling and neuronal differentiation pathways respectively. These findings suggest that hPSC-derived astrocytic progenitors retain similar molecular heterogeneity as that observed in human fetal brain and GBM tumor cells. Identification of specific astrocyte progenitors that exhibit distinct GBM subtype transcriptomic profiles should facilitate development of therapies that target specific GBM cell populations.

**Significance Statement:** Signaling pathways that create heterogeneity during development are essential for the generation of functionally distinct cell types that coordinate to achieve the complexity of the nervous system. However, studies have demonstrated that molecular dysfunction in stem/progenitor cells that give rise to the diverse cell types during normal development may contribute to malignant outcomes, including brain tumors such as glioblastoma. Using human pluripotent stem cell (hPSC) as a model, we found that developmental signals that promote the generation of astrocytes also give rise to astrocyte progenitor cells that share molecular similarities with different brain tumor subtypes. The study provides a novel hPSC differentiation protocol for generating molecularly distinct populations of astrocyte progenitors, which may be utilized in disease modeling or drug testing for specific brain tumor subtypes.

## Introduction

Glioblastoma multiforme (GBM) is a brain cancer known for its aggressive disease progression, post-treatment recurrence, and tumor heterogeneity (Ostrom et al., 2019). These properties have been attributed to the presence of tumorigenic glioma stem/progenitor cells (GSC), that display both proliferative and quiescent cell cycle states and that give rise to molecularly diverse progenies (Lathia et al., 2015). The GSC characteristics of GBM present challenges for existing disease treatments that provide only limited extension of patient lifespan (Piper et al., 2021). Better understanding of the molecular diversity and developmental lineage progression of GSC is essential for developing more effective therapeutics for GBM.

In the developing central nervous system, neural stem cells generated from a single layer of proliferating neuroepithelial cells undergo sequential neuronal then glial lineage specification stages through tightly controlled cellular and molecular mechanisms that dictate a series of hierarchical intermediate progenitor divisions with progressively limited proliferative and cell fate potential (Edlund and Jessell, 1999; Malatesta et al., 2003; Merkle et al., 2004; Kriegstein and Alvarez-Buylla, 2009; Huang et al., 2020). Many of the cell types that are involved in glial development are also present in brain tumors (Patel et al., 2014; Cuevas-Diaz Duran et al., 2019; Weng et al., 2019; Bhaduri et al., 2020; Couturier et al., 2020). Different populations of progenitor cells including neural progenitor cells (NPCs) and glial progenitor cells (GPCs) as well as astrocyte and oligodendrocyte precursors have been shown to be involved in GBM formation after mutation of molecules involved in regulating proliferation (e.g. EGFR or PDGFRA), of tumor suppressor genes (e.g. PTEN, TP53 or NF1), or other growth regulatory molecules (e.g. IDH1)(Ligon et al., 2007; Verhaak et al., 2010; Alcantara Llaguno et al., 2015; Venteicher et al., 2017). Notably, transcriptomic characterization of primary tumor samples suggests that there are at least three distinct subtypes of GBM, proneural, classical, and mesenchymal, that have different patterns of gene expression(Wang et al., 2017b; Teo et al., 2019), and some studies have suggested a fourth neural subtype (Verhaak et al., 2010) (NIH TCGA atlas). Recent single cell RNA sequencing of GBM tumors suggested the presence of heterogenous cell populations that share transcriptomic profiles of various developmentally identified glial progenitors (Tirosh et al., 2016; Venteicher et al., 2017; Neftel et al., 2019; Couturier et al., 2020). Understanding the mechanisms generating different intermediate progenitor cell types in the glial lineage during normal development is essential for deciphering the differences between normal and malignant cells, and more generally for devising regenerative strategies for repairing the damaged nervous system.

The emergence of human pluripotent stem cells (hPSCs) as a tool to study human brain development has advanced our understanding of the molecular mechanisms involved not only in normal development but also in disease (Espuny-Camacho et al., 2013; Lancaster et al., 2013; Duan et al., 2015; Chamling et al., 2021). Several protocols have been developed for the generation of astrocytes from hPSCs (Emdad et al., 2012; Mormone et al., 2014; Zhou et al., 2016; Sloan et al., 2017; Canals et al., 2018), but specific human astrocyte progenitors related to tumor subtypes have not been identified. To enable study of progenitor cell species that may be the origin cells for GBM, we developed a three stage, 160 day differentiation protocol for hPSCs that generates a heterogenous population of cells along the astrocytic lineage. Single cell RNA sequencing (scRNA-seq) identified two prospective astrocytic progenitor populations that share transcriptomic profiles with proneural or mesenchymal GBM subtypes. Using a combination of GBM-associated surface markers CD51, CD63, and CD71, we confirmed the presence of these astrocytic progenitors and determined that TGF-β signaling is the primary signaling pathway enriched in progenitors with mesenchymal properties. Taken together, our findings demonstrate the establishment of an hPSC-derived model *in vitro* that allows the identification, isolation and manipulation of GBM subtype-associated glial progenitor cells. Identification of progenitor populations that have been implicated in the recurrence of GBM after treatment should help to shape new therapies that specifically target these populations of cells.

## Materials and Methods

### hPSC Culture

Two iPSC lines were derived from normal human dermal fibroblasts and reprogrammed via non-integrating sendai virus as previously described (Seki et al., 2010). The H7 hESC line and the iPSCs were maintained on irradiated mouse embryonic fibroblasts in hESC medium containing Dublecco’s modified Eagle’s medium (DMEM)/F12, 20% knockout serum replacement, 0.1 mM nonessential amino acids, 1% Glutamax, and 0.1% 2-Mercaptoethanol. hESCs and iPSCs were expanded in feeder-free conditions on Matrigel (Corning) coated 6-well plates in mTeSR medium and passaged with ReLeSR (STEMCELL Technologies). iPSCs/hESCs were harvested as clumps with 1U/ml Dispase onto low adherent flasks in embryoid body (EB) medium containing Dulbecco’s modified Eagle’s medium (DMEM)/F12, 20% knockout serum replacement, 0.1mM nonessential amino acids, 2mM L-glutamine, 100μM β-mercaptoethanol without basic fibroblast growth factor (bFGF) to allow differentiation. Four days later, the medium was switched to N2 containing neurosphere medium (DMEM/F12, 0.1mM nonessential amino acids, 1X N2 supplement, 100μM β-mercaptoethanol, and 8mg/ml heparin) with 20ng/ml bFGF and 20ng/ml epidermal growth factor (EGF) for five days and changed every other day to maintain neural stem cells (NSCs) as neurospheres.

### Differentiation of hPSC-derived NSCs into Astrocytes

iPSCs/hESCs were differentiated into neural stem cells (NSCs) as outlined in supplemental methods. Neurospheres were manually picked under a dissection microscope and dissociated into single cells with Accutase (Millipore). 1.0×10^5^ cells were plated onto poly-L-ornithine (PLO)(15ug/ml; Sigma)/Laminin (10ug/ml; Roche) coated 24-well plates in differentiation medium containing DMEM (1x), DMEM/F12, B27 supplement, and 1ng/ml EGF. The medium was changed every 5 days. The cells were maintained in differentiation medium and passaged under the same plating density every 25 days or until confluent. At passage 2 the cells were treated with LIF (10ng/ml, R&D systems #7743-LF) for 7 days and cultured for an additional 7 days in basal media. Cells were then treated with recombinant human BMP4 (10ng/ml, R&D systems #314-BP) and followed by 7 days in basal media. Finally, cells were exposed to bFGF (10ng/ml, Thermo Fisher # 13256-029) for 7 days and followed by 7 days of basal media before harvested for analysis. For mature astrocyte differentiation, cells were passaged 25 days after bFGF treatments and maintained in astrocyte maturation media (AMM)(Zhang et al., 2016) for the remainder of the study.

### Immunocytochemistry

Cells cultured on glass coverslips were washed with 1x PBS, then fixed for 20 minutes at room temperature with cold 4% paraformaldehyde. Cells were then washed two times with 1x PBS and stored at 4 C until staining. Coverslips were incubated with blocking buffer (1% BSA, 0.25% Triton x-100, and 10% normal goat or donkey serum) for one hour at room temperature. Primary antibodies were diluted in staining buffer (1% BSA and 0.25% Triton x-100) and incubated overnight at 4 C. Cells were then washed 3 times with 1x PBS for 5 minutes each at room temperature. AlexaFluor secondary antibodies (Thermo Fisher) were diluted in staining buffer at 1:2000 and cells were incubated for one hour at room temperature, Hoechst was used as a nuclear stain. After washing 3 times with 1x PBS for 5 minutes each, coverslips were mounted with Prolong Gold. Primary antibodies used in this study are as follow: Acsbg1 (Abcam #ab65154, IHC at 1:100), CD51 (Biolegend #327910, FACS at 1:500), CD51 (Cell Signaling Technology #13208 1CC at 1:1000), CD63 (Millipore Sigma #C7930, FACS and ICC at 1:500), CD71 (Biolegend #334208, FACS and ICC at 1:1000), Doublcortin (Novus Bio. # NBP1-92684, ICC at 1:250), GJA1 (Santa Cruz Biotech. #sc271837, ICC at 1:500), GFAP (DAKO #GA52461-2, ICC at 1:1000), Ki67 (Abcam #ab16667, ICC at 1:1000), Nestin (Abcam #ab18102 ICC at 1:1000), S100b (Millipore Sigma #S2532, ICC at 1:1000), SPARCL (Thermo Fisher #pa5-47375, ICC at 1:500), Tuj1 (Millipore Sigma #T8660, ICC at 1:100).

### Image acquisition and statistical analysis

Images obtained for the differentiation experiments were taken on a confocal microscope (Leica SP5) at 20x and 40x and exported to NIH image J for cell number quantification. Data analysis was conducted in Prism 9 (Graphpad) using the statistical tests specified by each experiment.

### FACS

Accutase (Millipore) dissociated cells were resuspended in 100ul of cell staining buffer and incubated with 5ul of Human TruStain FcX blocking solution (BioLegend 422303) for 10 minutes at room temperature. After blocking, 5ul of PE anti-human CD51 (BioLegend 327910), APC anti-human CD71 (BioLegend 334108), and Brilliant Violet 421 anti-human CD63 (BioLegend 353029) antibodies were added to the cells and incubated for 20 minutes on ice and washed three times with cell staining buffer with intermittent pelleting at 900 rpm for 5 minutes. The samples were resuspended in 400ul of cell staining buffer and incubated on ice until sorted on a FACSAria II (BD Biosciences). To determine the auto fluorescence of the cell, an unstained control was included as well as samples incubated with CD51 only, CD71 only, and CD63 only.

### Glutamate Assay

Cells were incubated in Hank’s balanced salt solution (HBSS) buffer (Corning) for 30mins. The cells were then incubated in HBSS containing 50 uM glutamate (Sigma) for 1 hour in a 37 C degree incubator. To determine the amount of glutamate uptake, HBSS containing 50uM of glutamate was added to empty wells and incubated for 1 hour. Medium was collected from each well after 1 hour and analyzed with a colorimetric glutamate assay kit (Sigma MAK004-1KT), according to the manufacturer’s instructions.

### Calcium Imaging

hPSC-derived astrocytes were cultured on glass-bottom fluorodishes (FD35-100) coated with poly-L-ornithine (PLO)(15ug/ml; Sigma)/Laminin (10ug/ml; Roche) at a density of 20,000 cells/dish. Calcium imaging was performed using Fura-2 AM (Invitrogen F1221) fluorescent dye. Samples were loaded with 1xACSF containing 2μM Fura-2 for 30 minutes at 37°C and subsequently washed 3x in 1x ACSF. Cells were allowed to de-esterify for 30 min at RT prior to visualization. Cells were imaged on Nikon Diaphot inverted epifluorescence microscope. Fluorescence was monitored at 520 nm after excitation at 340 nm (bound Ca2+) and 380 nm (free Ca2+) and images were acquired every 6s. Regions of interest (ROIs) were made around the cell soma and fluorescence intensity was measured for each image over a period of 500s. The F340/F380 ratios were calculated for each time point. Ratios of F340/F380 were collected before and during treatment with ATP (50µM) using MetaFluor software from Universal Imaging Corporation (West Chester, PA).

### RNA Isolation and qRT-PCR

RNA was collected from cells at the NPC, LBF and AMM stage with the RNAqueous Micro total RNA isolation kit (Thermo Fisher AM1931) according to the manufacturer’s instructions. cDNA was synthesized using the SuperScript IV VILO Master Mix kit (Thermo Fisher 11766500), and qPCR conducted using the SYBR Green PCR Master Mix (Thermo Fisher 4312704). DNA sequence for primers used in qPCR analyses are listed in extended data table 1-1.

### Single Cell RNA Sequencing

Cultured astrocyte progenitors differentiated from hESCs at 160 and 220 DIV were dissociated and processed for single cell RNA sequencing with the Chromium Single Cell 3’ Solution V3.1 (10x Genomics) with assistance from the University of Chicago genomics core. For the day 160 sample, 90,000 cells from three independent differentiations are combined, captured and barcoded on microfluidic chip, subsequently used for cDNA library construction before sequencing on the Illumina Hiseq 4000. Cell Ranger (version 7) was used to align the FASTQ files for each sample to the human reference genome (hg38) and count the number of reads from each cell that align to each gene. The matrix files which summarize the alignment results and were then imported in Seurat (v4, Satija Lab, NYGC) for further analysis. The Seurat object data was first filtered by total count and mitochondrial DNA content to eliminate doublets and poor quality cells respectively, followed by normalization and scaling using R package *SCTransform* (Hafemeister and Satija, 2019) with cell cycle genes regressed as described (Tirosh et al., 2016). Cell clusters were identified using snn resolution of 0.5, followed by identification of cluster specific DEG using Seurat function *FindAllMarkers* with logfc and min.pct thresholds of 0.25. Cell cluster distribution in two dimensions were visualization using uniform manifold approximation and projection (UMAP) dimensional reduction, and expression of genes of interest on the UMAP was presented as using the *FeaturePlot* function in Seurat. To recapitulate FACS gating thresholds, identification of CD51, CD63, and CD71 expressing cells in the scRNA-seq dataset was performed with the Seurat *WhichCells* function with threshold set at 1.5. *Monocle* (v3) R package was used for lineage trajectory analysis with assistance from the University of Chicago CRI bioinformatics core, RRID:SCR_022937. Pathway analysis was performed on the NCATS BioPlanet (2019) platform(Huang et al., 2019) using Enrichr (Kuleshov et al., 2016). Raw and matrix data from the scRNA-seq is deposited at NCBI Gene Expression Omnibus repository at accession number GSE226740.

### Bulk RNA Sequencing Analysis

Cultured astrocyte progenitors at 160 DIV generated from hESCswere dissociated and segregated into 4 populations via FACS as previously described. RNA was extracted from isolated cells using the RNAqueous Micro total RNA isolation kit (Thermo Fisher AM1931) according to manufacturer’s instructions. The university of Chicago Center of Research Informatics’s (CRI) pipeline for Illumina RNA-seq data was utilized for data analysis. The raw RNA-seq data went through quality control assessment at first. After reviewing the QC stats, sequence alignment, expression quantification and normalization were performed. Further, tertiary analysis, such as significant differential expressed gene identification, and pathway (IPA) enrichment analysis were conducted. The quality of raw sequencing data was assessed using fastQC v0.11.5 and multiQC v1.7. Reads were first went through quality trimming using Trimmomatic v0.36(Bolger et al., 2014). The remaining reads were mapped to the human reference genome (hg38)(Frankish et al., 2019) using STAR 2.7.2a(Dobin et al., 2013). Reads were further quantified to genomic features using featureCounts v1.5.3 (Liao et al., 2014) in Subread package(Liao et al., 2013) with the gene annotation included in GENCODE Release 31. Sample-based quantified read counts of 12 samples were merged into count table, and genes with low expression (the sum of counts across all samples is less than 1) were removed for further downstream differential analysis. Normalized gene expression values were obtained through TMM normalization (Robinson and Oshlack, 2010)with library size correction. Expression normalization and differential analysis were both performed using R bioconductor package edgeR (Robinson et al., 2010; McCarthy et al., 2012). Raw transcript count data is deposited at GEO repository at accession number GSE226647.

## Results

### Characterization of multipotential progenitors and astrocytes derived from hPSCs

Progress in the differentiation of hPSCs into astrocytes has resulted in protocols that take less time than the normal process of lineage commitment *in vivo,* and most aim to generate post-mitotic astrocytes and are not designed to preserve intermediate glial progenitor cell species. Truncation of the process of astroglial differentiation potentially limits the ability to identify intermediate progenitor species that share molecular similarities with subtypes of cells detected in GBM. We therefore adapted our previously published protocol for differentiating hPSCs into astrocytes(Duan et al., 2015) to maximize the number of astroglial progenitor species in the cultures. hPSC derived neural progenitor cells (NPCs) were dissociated into single cells and maintained as adherent cultures that spontaneously differentiated over 3 passages for 100 days *in vitro* (DIV) before sequential treatment with leukemia inhibitory factor (LIF), bone morphogenetic protein 4 (BMP4), and basic fibroblast growth factor (bFGF) (Fig 1A). Immunocytochemical (ICC) analyses revealed that addition of LIF significantly increased the number of GFAP^+^ and S100β^+^ cells compared to untreated controls (Fig 1B-C). The majority of GFAP^+^ cells at this stage co-expressed Nestin, suggesting that the GFAP-expressing cells remained as progenitor cells after LIF treatment, similar to findings in mouse (Bonaguidi et al., 2005). To promote further astrocytic lineage commitment and differentiation, LIF-induced cells were treated with BMP4 at 125 DIV. BMP4 treatment increased the number of cells expressing GFAP and S100β and increased the size and number of processes of the cells. Simultaneously, there was a decrease in expression of Nestin, Tuj1, and Ki67 (Fig 1B-C), similar to BMP4 effects on mouse astrocyte lineage commitment(Gross et al., 1996; Bonaguidi et al., 2005; Scholze et al., 2014). Cells were passaged following BMP4 treatment and treated with bFGF at 146 DIV to keep the cells in the cell cycle and to delay maturation of the glial progenitor cell species. ICC analyses at 160 DIV showed an increase in the number of S100β+ cells and a decrease in the number of GFAP+ cells, consistent with reported findings(Roybon et al., 2013; Savchenko et al., 2019). However, there also was a small number of Nestin and Tuj1 positive cells, suggesting the persistence of some neurogenic NPCs. These findings suggest that sequential treatment with LIF, BMP4, and bFGF (hereafter referred to as LBF) was sufficient to generate primarily gliogenic NPCs and astrocytes, although the presence of some neuronal lineage cells was also detected. LBF stage cells can further be guided toward astrocytic differentiation and maturation when cultured in previously published astrocyte maturation media (AMM)(Zhang et al., 2016), and maintained up to 250 DIV. Glutamate uptake and calcium imaging assays, as well as single cell RNA sequencing analysis of cells cultured at the 220 DIV timepoint, confirmed that the majority of AMM cells share functional and transcriptomic similarities with mature astrocytes (Extended data fig. 1-1).

**Figure 1.**
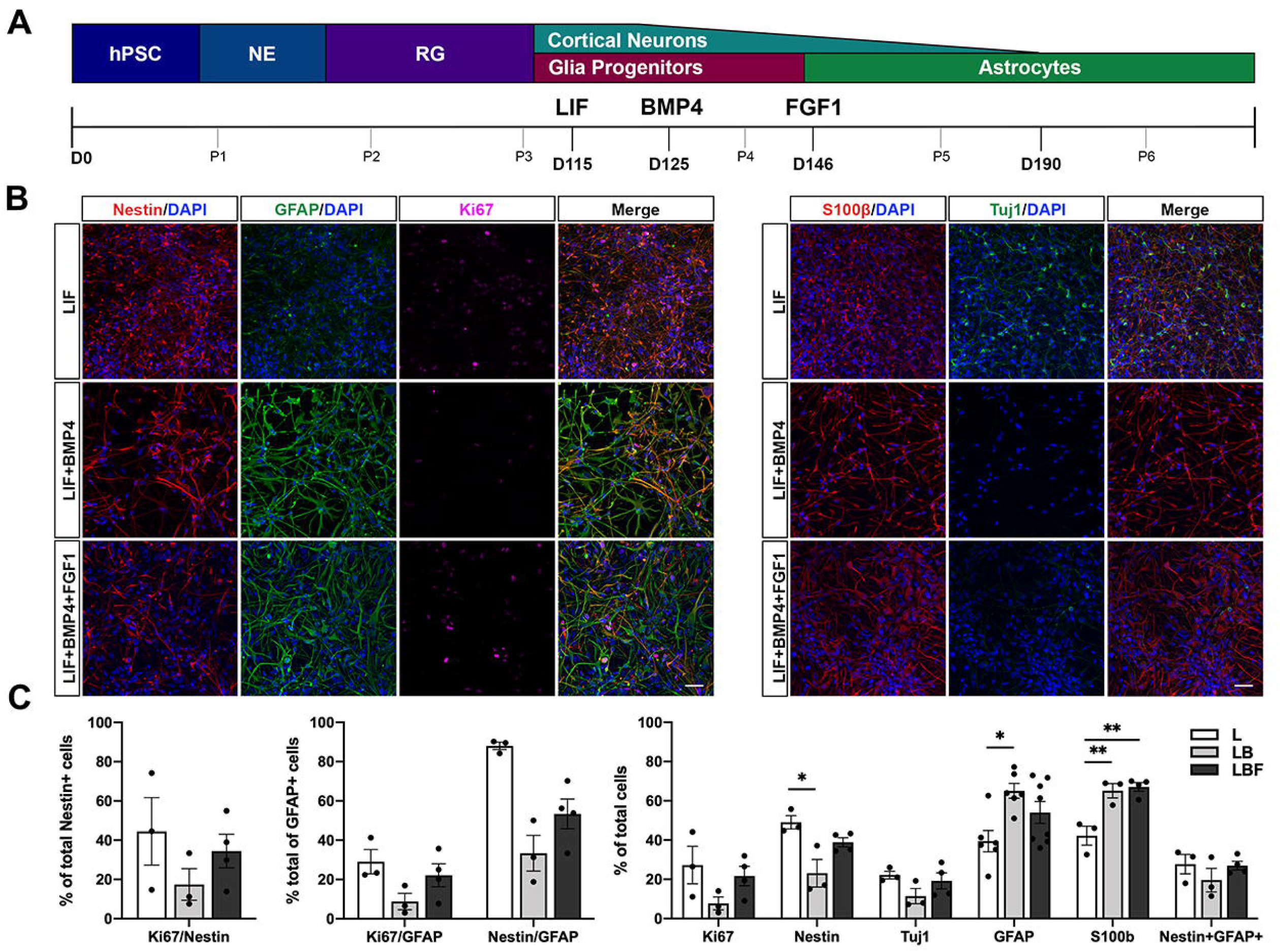
Generation of distinct astrocytic progenitor subtypes from hPSCs. (**A**) Experimental timeline of the astrocyte progenitor differentiation protocol. (**B**) Immunostaining of GFAP, S100β, Nestin and Tuj1 at different time points of the differentiation protocol, including after the addition of LIF (L, *n*=3), after sequential addition of LIF and BMP4 (LIF+BMP4; LB, *n*=4), and after sequential addition of LIF, BMP4, and bFGF (LIF+BMP4+bFGF; LBF, *n*=6). (**C**) Quantification of KI67+, Nestin+, Tuj1+, GFAP+ and S100β+ cells to total number of DAPI+ nuclei for the three timepoints. *n* represents the number of independent differentiation experiments from one ESC line and two iPSC lines. Statistical significance measured by one-way ANOVA followed by Tukey’s post-hoc test, *p<0.05, **p<0.01.

### Single Cell Transcriptomic Profiling of LBF-induced astrocytic progenitors identified distinct populations correlating with GBM subtypes

We next performed single cell RNA sequencing (scRNA-seq) to unbiasedly characterize the cellular heterogeneity of the hPSC-derived progenitors. Unsupervised clustering of cellular populations based on similarity in overall transcriptomic profiles revealed ten different clusters (Fig 2A, Extended data table 2-1). Clusters Ast.1, Ast.2 and Ast.3 contained astrocytic progenitors with varying expression of typical astrocyte markers (GFAP, S100β, HTRA1, SPARCL1, Fig 2B, 2C). To classify the 3 different astrocytic lineage committed cell clusters, we examined expression of a collection of mature and fetal human astrocyte enriched genes (Zhang et al., 2016) (Extended data table 2-2) and found that mature astrocyte enriched genes are most highly expressed in cluster Ast.1 and least expressed in cluster Ast.2 (Extended data fig. 2-1A). On the other hand, fetal astrocyte enriched genes are more highly expressed in clusters Int.1, Int.2, and Stem/Mitotic, which are also enriched in genes related to cell cycle (MKi67, CENPF, Fig 2B, 2C) thus indicating they are cycling stem or intermediate neural progenitors. Cluster Neu.1, Neu.2, and Neu.3 contained progenitors that are express molecular markers of to the neuronal lineage, with high expression of DCX, CALB2 (Fig 2B, 2C). SOX9, which is essential for astrocytic lineage commitment and maintenance of the progenitor fate (Stolt et al., 2003; Scott et al., 2010; Sun et al., 2017), was expressed in all clusters. None of the clusters expressed significant levels of genes associated with postmitotic neurons (SYN1, Fig 2C) or oligodendrocytes (MBP1, Fig 2C), suggesting that cultures consisted of lineage restricted astroglial and neuronal progenitors that have not yet fully differentiated.

**Figure 2.**
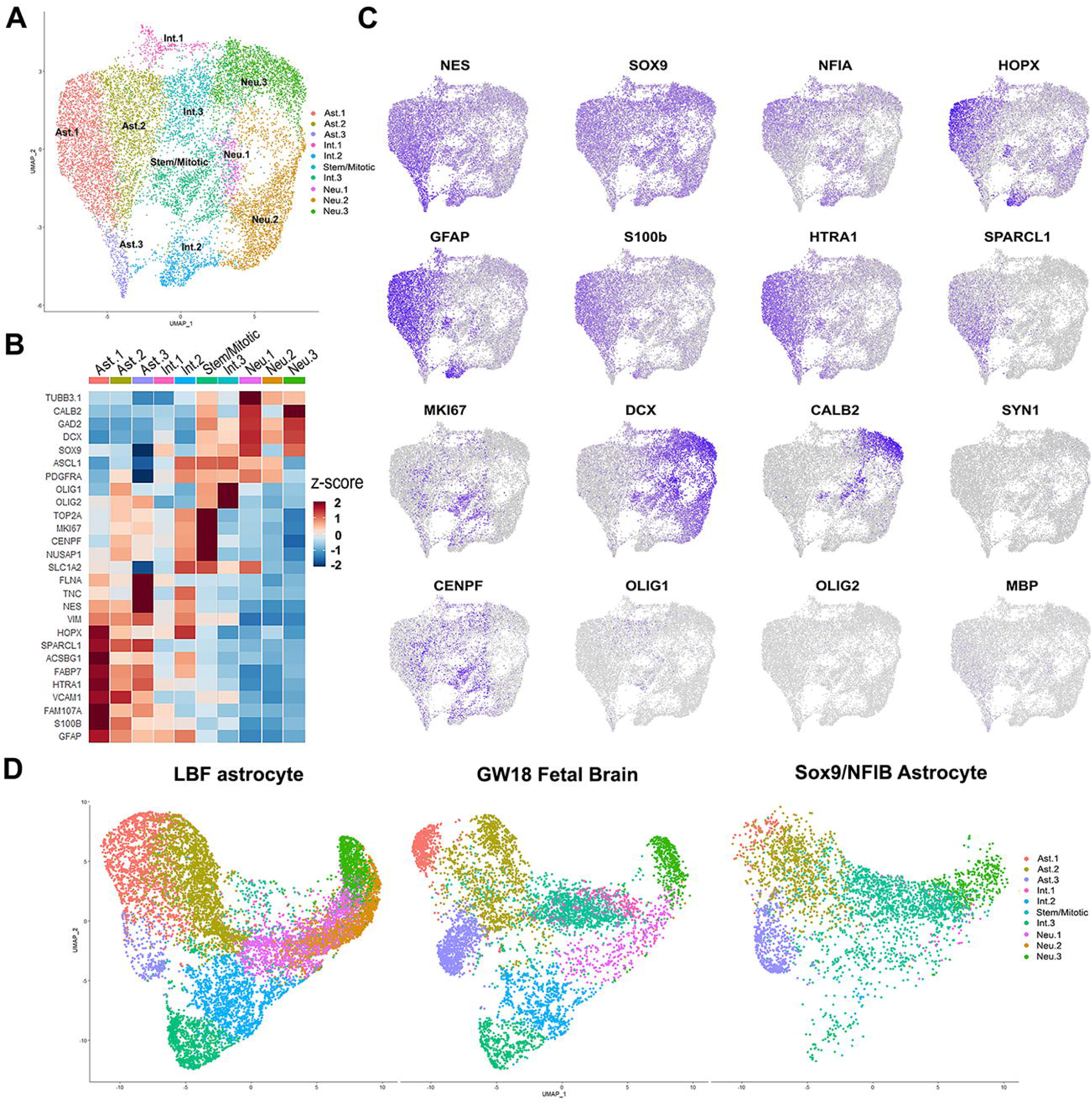
Single cell RNA sequencing analysis demonstrate transcriptomic heterogeneity in hPSC-derived astrocyte progenitors. (**A**) Uniform manifold approximation and projection (UMAP) plot of scRNA-seq data from hPSC-derived astrocyte progenitors at the LBF stage (*n*=11,817). In total, 10 clusters were identified. (**B**) Heatmap representation of neural cell lineage specific genes confirmed the presence of multiple neural progenitors, intermediate progenitor, astrocytic and neuronal populations among cell clusters identified in the scRNA-seq. (**C**) UMAP feature plots of neural progenitor enriched genes (NES, SOX9, NF1A, HOPX), astrocyte enriched genes (GFAP, S100β, HtrA1, and SPARCL1), cell-cycle genes (MKI67, CENF), and neuronal genes (DCX, CALB2, and SYN1), illustrating the spatial distribution of lineage segregated cell clusters. Specifically, astrocyte committed clusters on the left, neuronal committed clusters on the right, and cycling progenitors near the center of the UMAP plot. (**D**) Integrated UMAP plot of scRNA-seq dataset comparisons between LBF-induced cells, primary cells from gestation week 18 human fetal neocortex, and hPSC-derived transcription factor induced-astrocytes. The distribution of aligned cell clusters across the three samples demonstrated greater similarities in the number of cell clusters identified in LBF-induced cells and primary brain tissue. *n* represents the combined number of cells collected from three independent differentiation experiments from one embryonic stem cell line. Data are represented as means ± s.e.m. Scale bar, 50µm.

To compare how cell populations identified in the scRNA-seq align with progenitors present during normal human development, we performed meta-analysis using published scRNA-seq datasets of human neocortex isolated from gestation week 18 (GW18) fetal tissue (Barbar et al., 2020). Integration and alignment of the two datasets revealed significant transcriptomic and cell type similarities between LBF-induced cells and primary cells (Fig 2D), albeit with clear variations in the percentage of cells represented in each cluster between the experimental samples. This finding suggests that LBF-induction, at least in part, recapitulates the generation of heterogeneous intermediate progenitors observed during *in vivo* development. In contrast, astrocytes generated using viral transduction of SOX9 and NFIB(Canals et al., 2018) showed similarities with LBF-induced cells in the astrocyte-committed Ast.1-3 progenitor populations, but significantly reduced numbers of dividing progenitors (Stem, Int.1-2) and neuronal-committed (Neu.1-2) populations that are detected in both GW18 fetal brain tissue and cells from LBF induction (Fig 2D). Taken together, these findings further confirmed that the LBF induction protocol generates heterogenous, multi-lineage neural progenitors similar to that observed during human cortical gliogenesis.

### Identification of glial progenitor populations that share transcriptomic profiles with GBM subtypes

It has recently been reported that glioma tumor composition reflects the heterogeneous nature of glial lineage progression observed during normal neural developmental(Patel et al., 2014; Tirosh et al., 2016; Filbin et al., 2018; Couturier et al., 2020; Hamed et al., 2022). We asked whether cell populations identified after LBF-induction could share transcriptomic similarities with cells detected in tumors of different GBM subtypes. We first examined relative average expression of GBM subtype enriched genes from mesenchymal, proneural, and classical GBMs in all cell clusters identified in our scRNA-seq analysis (Fig 3A). Mesenchymal GBM associated genes were highly enriched in cluster Ast.3, whereas proneural GBM associated genes were enriched in cluster Neu.1 (Fig 3A). Markers of classical and neural GBM were detected at higher levels in cluster Ast.1, although not as clearly enriched over cluster Neu.1 (Fig 3A, Extended data fig 2-1B). Comparison of GBM subtype markers from another reported dataset(Wang et al., 2017a) also revealed similar correlations between cluster Ast.3 and mesenchymal GBM, as well as cluster Neu.1 and proneural GBM (Extended data fig. 2-1C). Feature plot expression analyses of select mesenchymal GBM markers SERPINE1, TAGLN, and ACTA2 also demonstrated enrichments in cluster Ast.3, supporting the expression heatmap results (Fig 3B). Similar analyses with proneural GBM markers DLL3, SCG3, and NCAM1 revealed that cluster Neu.1 expresses all three markers although not exclusively. These findings confirm the presence of molecularly distinct progenitors in the cultures that share similar expression profiles with cells detected in mesenchymal and proneural GBM.

**Figure 3.**
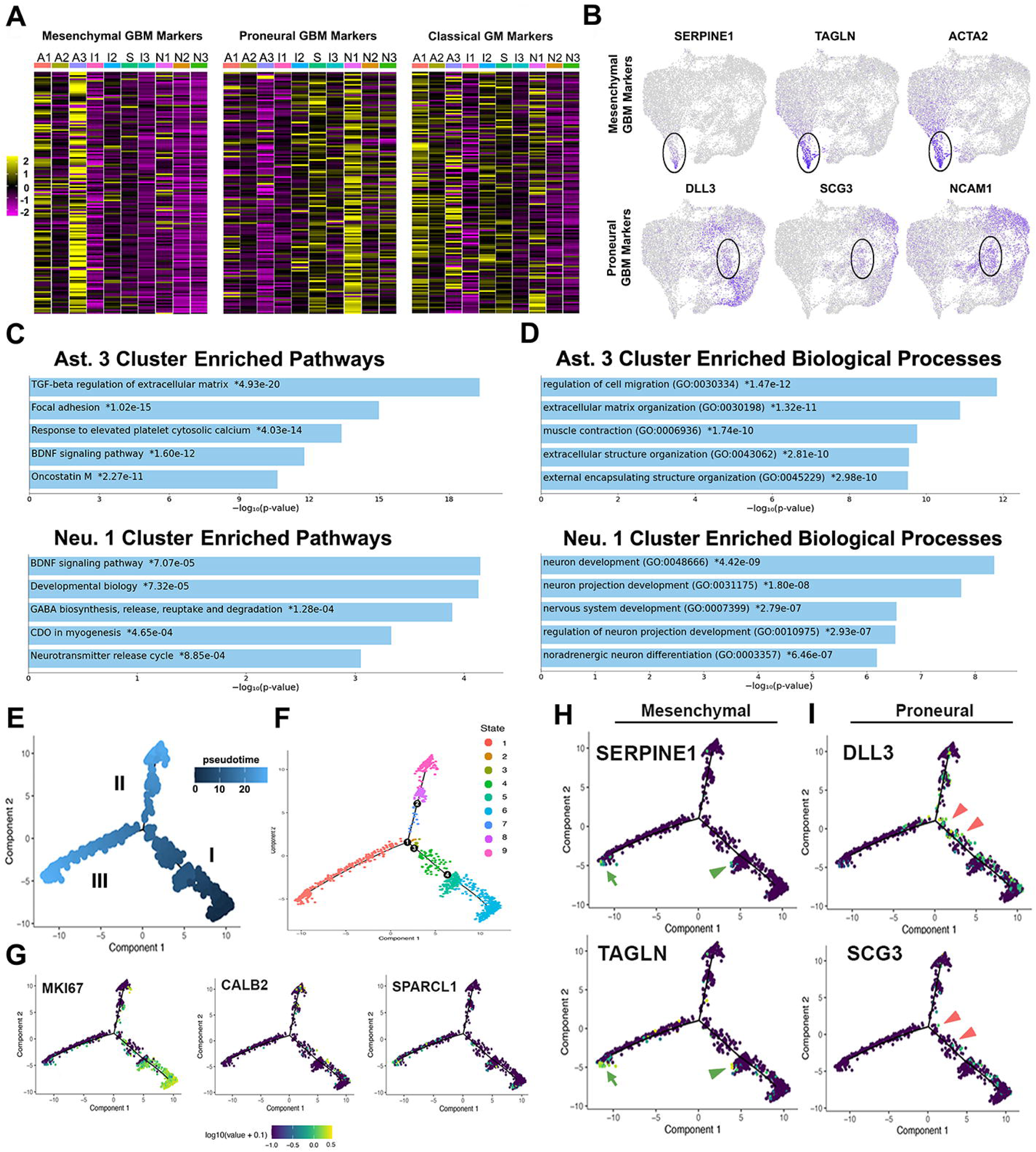
Transcriptomic analysis revealed correlation between gene expression profiles of hPSC-derived astrocyte progenitor populations and GBM subtypes. (**A**) Heatmap representation of relative expression of genes enriched in mesenchymal, proneural and classical GBM subtypes across the 10 identified clusters. High expression of mesenchymal GBM-associated genes is detected in astrocytic cluster Ast.3 (A3), and high expression of proneural GBM-associated genes is detected in neuronal cluster Neu.1 (N1) (**B**) Feature plots of select mesenchymal GBM-associated genes (SERPINE1, TAGLN, ACTA2) and proneural GBM-associated genes (DLL3, SCG3, NCAM1). (**C**) Top 5 results of Pathway analysis of DEG enriched in cluster Ast.3 and Neu.1 (**D**) Top 5 results of Gene ontology analysis of biological processes associated with DEG enriched in cluster Ast.3 and Neu.1. (**E**) Pseudotime trajectory plot based on scRNA-seq data showing two branching differentiation paths from cycling neural progenitors. (**F**) Cells along the pseudotime trajectory are divided into 9 different states. States 2-6 are neural progenitor cells with differences in cell-cycle state. States 7 and 8 are progenitor cells along the neuronal lineage, state 9 cells are immature neurons, and state 1 cells are astrocytes. (**G**) Distribution of cell type-specific markers for progenitors (MKI67), neurons (CALB2), and astrocytes (SPARCL1) along the pseudotime trajectory. (**H, I**) Distribution of cells expressing mesenchymal (SERPINE1, TAGLN) or proneural (DLL3, SCG3) GBM subtype markers along the pseudotime trajectory.

To further explore the molecular heterogeneity between cell clusters identified in the single cell RNA-seq, we selected differentially expressed genes (DEG) in each cluster that are increased greater than 1.5-fold over all remaining clusters and performed pathway analyses with a focus on clusters Ast.3 and Neu.1. We found that TGF-β signaling regulation of extracellular matrix as the top pathway enriched in cluster Ast.3, whereas BDNF signaling was identified as the top pathway enriched for cluster Neu1 (Fig 3C). Gene ontology analysis of biological processes associated with cluster Ast.3 DEG revealed enrichments in regulation of cell migration and extracellular matrix organization, consistent with the role of TGF-β signaling in epithelial to mesenchymal transition (Fig 3D). The same analysis with Neu.1 DEG showed neuronal development as the top enriched biological process (Fig 3D). Taken together, these findings further suggest that clusters Ast.3 and Neu.1 share gene expression profiles with mesenchymal and proneural GBM respectively.

### Cell lineage trajectory analysis further demonstrates progenitor bipotentiality

To investigate how GBM-associated progenitors are related to other cell populations in our model system, we asked whether the lineal relationships between cell clusters identified in the scRNA-seq can be inferred using cell trajectory reconstruction algorithms Monocle (Trapnell et al., 2014). Analysis of the pseudotemporal trajectory demonstrated two paths of lineage specification of NPCs located in path I as they differentiate and mature (Fig 3E). The analysis also revealed 9 distinct cellular states, which we mapped to the cell clusters identified in Seurat based on corresponding DEG between the two analyses (Fig 3F). Cells in states 3-6 are NPCs with differences in cell-cycle state, highlighted by the expression MKi67 (Fig 3G). Cells in states 7-9 are cells committed along the neuronal lineage, as indicated by the enrichment of CALB2 toward the distal end of the trajectory path (Fig 3G). Cells in state 1 include cells committed to the astrocytic lineage, demonstrated by the enriched expression of SPARCL1 along the trajectory path (Fig 3G).

We next examined expression of mesenchymal and proneural GBM enriched genes along the pseudotemporal trajectory and found that mesenchymal GBM subtype enriched genes SERPINE1 and TAGLN are mostly located at the distal end of path III, corresponding to astrocytes with more differentiated expression profiles (Fig 3H, green arrow). When detected in path I, the expressed cells are located along the left half of the cell clusters, suggesting an astrocytic lineage bias (Fig 3H, green arrowheads). Proneural GBM subtype enriched genes DLL3 and SCG3 appear along right half of the path I (Fig 3I, red arrowheads), continue into committed path II but excluded at the most distal tip, suggesting that proneural GBM markers are expressed in multiple intermediate progenitor populations during neuronal specification, but less so in more differentiated neuroblasts. Interestingly, cells with mesenchymal marker expression at the most distal tip of path III also express MKi67, suggesting that proliferation in mature astrocyte may be correlated with acquisition of mesenchymal properties (Fig 3G). This analysis further suggests that bipotential progenitors present in the culture are lineally related to more committed cell types observed at this point of development, and that cell populations associated with mesenchymal and proneural GBM subtypes are positioned at distinct points along the pseudo-temporal timeline.

### Cell surface markers identify distinct cell populations at the LBF stage

To understand how heterogeneity of intermediate progenitors may contribute to the cellular diversity in GBM, we sought to identify and isolate different populations of NPCs and their progenies in the hPSC-derived cultures. In mice, expression of the cell surface markers CD51, CD63, and CD71 correlates with distinct mouse astrocytes and tumor subtypes (John Lin et al., 2017). We therefore determined whether the combinatorial expression of these surface markers could identify distinct human progenitor populations that correlate with different human GBM subtypes. ICC at 160 DIV for CD51, CD63, and CD71 revealed cell populations identified by differential and combinatorial expression of these surface markers (Fig 4A-4B). Co-expression analysis of CD51, CD63 and GFAP revealed that 67% of GFAP+ cells are CD51+CD63+ and 38.1% of CD51+CD63+ are GFAP+ (Fig 4C, 4D), correlating CD51 and CD63 co-expression with astrocytic fate. A similar distribution of surface markers was detected using fluorescence activated cell sorting (FACS, Fig. 4E, 4F). Based on combinatorial expression of the three surface markers, we were able to identify and FACS-isolate four distinct CD51-expressing populations (CD51+CD71+CD63+, CD51+CD71-CD63-, CD51+CD71+CD63- and CD51+CD71-CD63+; hereafter referred to as population 1, 2, 3, and 4 respectively) (Fig 4G). Quantification of cell number in each population revealed that the triple positive population 1 was the predominant population (44.8%) across multiple differentiation experiments (Fig 4F), followed by the CD51 only population 2 (9.1%, Fig. 4F). Populations 3 and 4 together comprised 15.9% of the sorted cells (Fig 4F).

**Figure 4.**
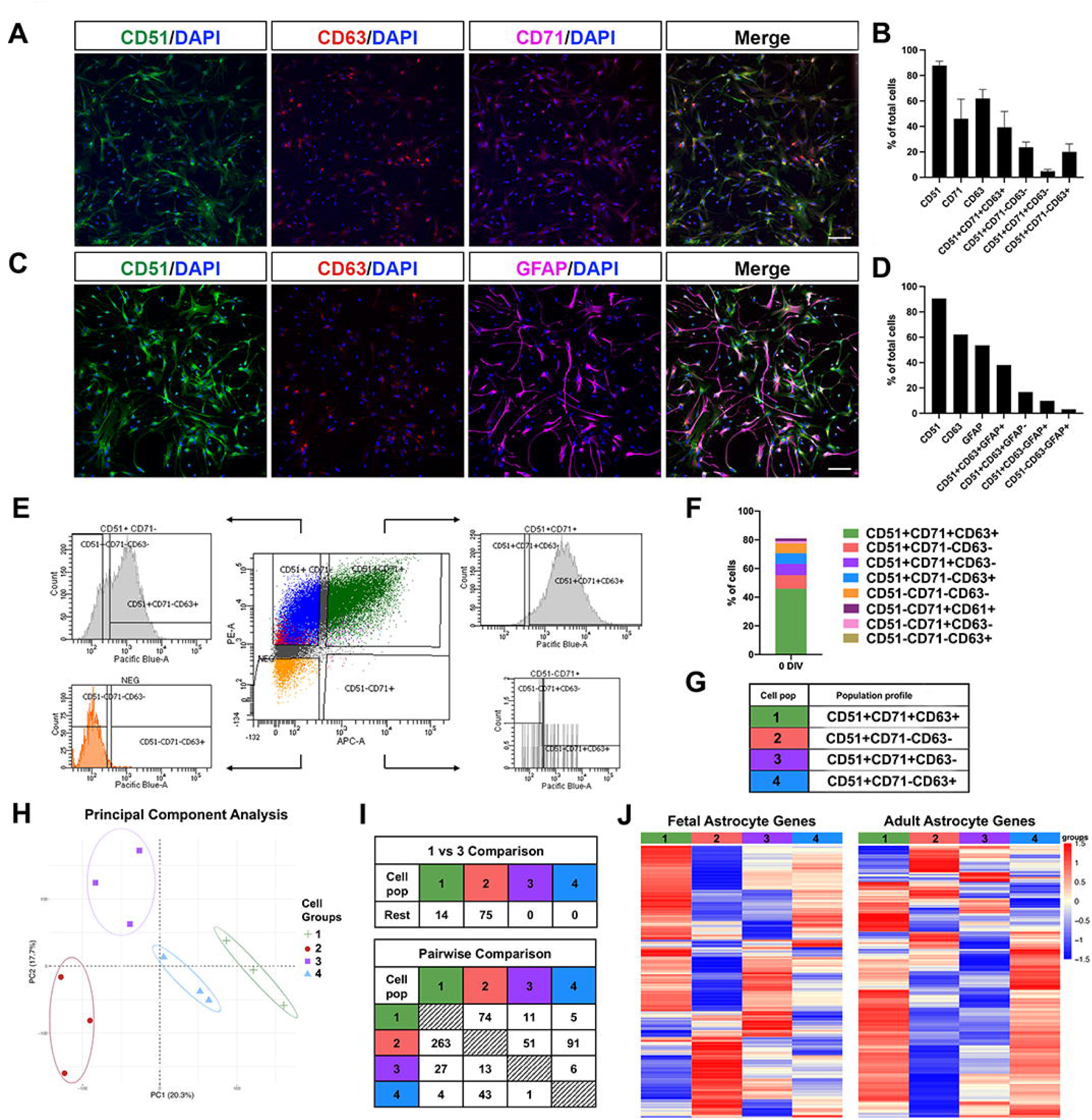
Combinatorial expression of CD51, CD63, and CD71 identify four distinct populations present during the mid-phase of astrocyte differentiation. (**A**) Immunostaining of CD51, CD63 and CD71 in LBF treated cells at 160 DIV (*n*=3). (**B**) Quantification of the percentage of LBF stage cells with CD51, CD63, and CD71 immunostaining and different combinations of co-expression (*n*=3). (**C**) Immunostaining of CD51, CD63 and GFAP in LBF treated cells (*n*=3). (**D**) Quantification of the percentage of LBF stage cells with CD51, CD63 and GFAP immunostaining and different combination of co-expression (*n*=3). (**E**) Histogram of flow cytometry gating strategy for sorting CD51, CD71, and CD63 expressing or co-expressing cells after LBF treatment. (**F**) Proportion of FACS-isolated cells with each possible combination of CD51, CD71, and CD63 co-expression (*n*=6). (**G**) Labeling of the four predominant populations identified based on CD51, CD71, and CD63 expression. Population 1 (CD51+CD71+CD63+, green), population 2 (CD51+CD71-CD63-, red), population 3 (CD51+CD71+CD63-, purple), and population 4 (CD51+CD71-CD63+, blue) account for 70% of FACS-isolated cells. (**H**) Principal component analysis (PCA) plot illustrating the overall sample distribution based on transcriptomic profiles of populations 1-4 cells used for RNA-seq. (**I**) Table depicting the number of differentially expressed genes from pairwise and 3 versus 1 comparison analyses based on greater than 1.5-fold difference and false discovery rate of 0.05. (**J**) Heatmap representation of relative expression of fetal and mature astrocyte enriched genes in each population. *n* represents the number of independent differentiation experiments from one ESC line and two iPSC lines. Scale bar, 50µm.

We next performed RNA sequencing to delineate the molecular profiles of these four populations (supplementary table 2). Principal component analysis revealed that each population, from three independent differentiation experiments, clustered together indicating that each population has a conserved molecular signature (Fig 4H). To further define the similarity or differences among the groups, we quantified the number of DEG from each group in both 1 vs 3 and pairwise comparisons (Fig 4I, Extended data table 4-1). These analyses demonstrated that populations 1 and 2 have more divergent transcriptomes compared to differences between populations 1 and 3 or 1 and 4 (Fig 4I). These findings confirmed the molecular heterogeneity among the 4 populations and highlight the differences between populations 1 and 2.

To determine the developmental stage of the 4 progenitor populations along the astrocytic differentiation timeline, we examined expression of genes enriched for human fetal or adult astrocytes (Zhang et al., 2016). Fetal astrocyte enriched genes related to astrocytic fate are most prevalent in population 1, whereas fetal astrocyte enriched genes related to progenitor fate are highly expressed in population 2 (Fig 4J). Mature astrocyte genes are preferentially detected in population 1 and 4, and least detected in population 2 (Fig 4J). Taken together, our combinatorial surface marker approach identified astrocytic progenitor populations 1 and 2 as two clearly molecularly distinct cellular populations that arise during our LBF-induced astrocyte differentiation protocol.

### Correlation of hPSC-derived astrocyte progenitors gene expression profiles with GBM subtypes

We next examined expression of GBM subtype enriched genes in the surface marker isolated cell populations. Mesenchymal GBM subtype enriched genes identified in two different studies (Verhaak et al., 2010; Teo et al., 2019), are highly expressed in population 1 and detected at much lower levels in population 2-4 (Fig 5A, 5B). The majority of proneural GBM subtype enriched genes are highly expressed in population 2 and are found at much lower levels of expression in population 1, 3 and 4 (Fig 5A). These findings suggest that combinatorial analysis of CD51, CD63, and CD71 expression in our hPSC-derived cultures identifies astrocytic progenitor populations that share gene expression profiles with mesenchymal and proneural GBM subtypes.

**Figure 5.**
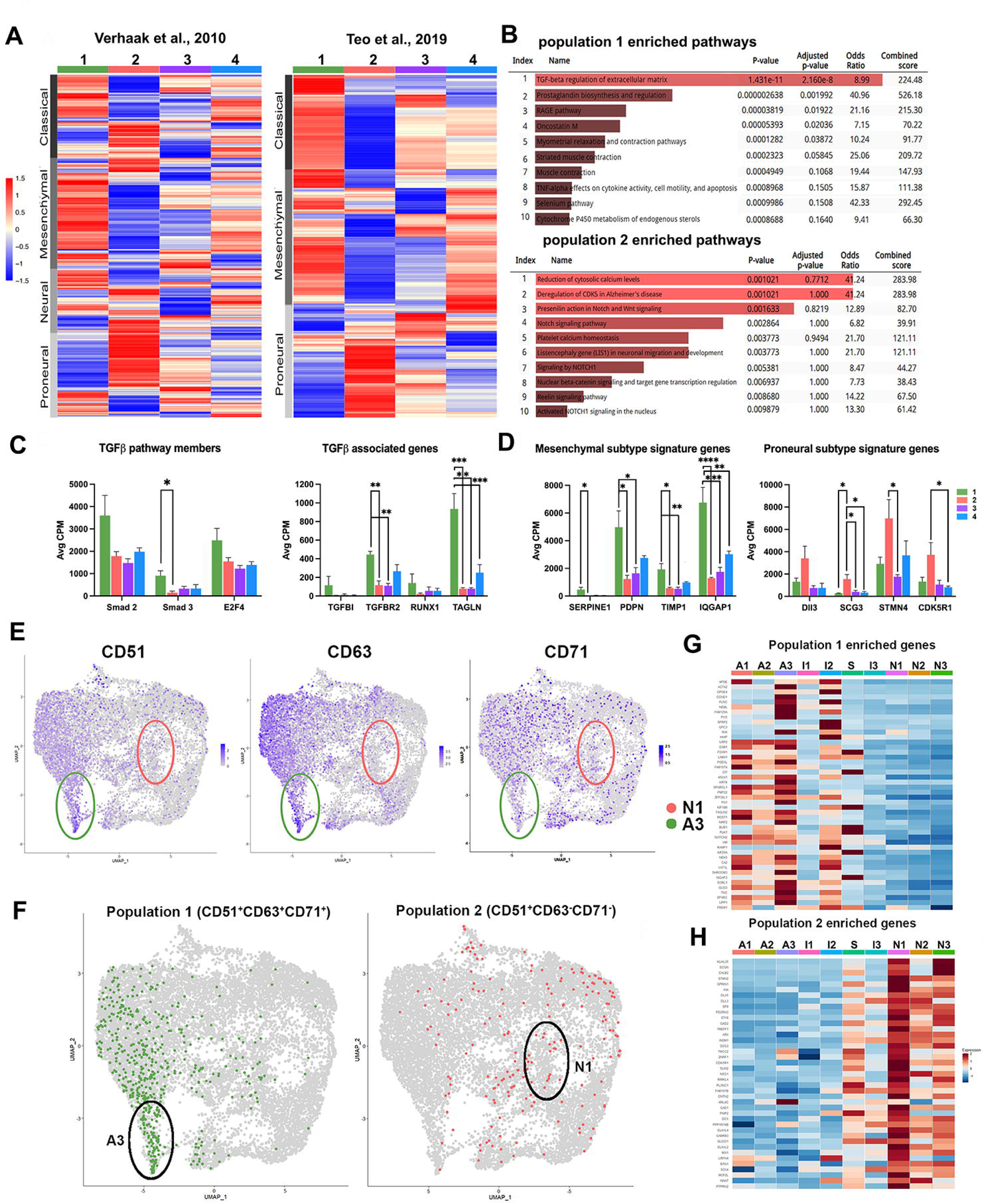
Surface markers identified populations 1 and 2 share gene expression profiles with mesenchymal and proneural GBM subtypes respectively. (**A**) Heatmap representation of relative expression of GBM subtype-enriched genes in FACS isolated populations 1-4 cells GBM subtype enriched gene list obtained from Verhaak et al (2010). (**B**) Pathway analyses of differentially expressed genes identified in populations 1 and 2 highlighted enrichment in TGF-β signaling in population 1 and Notch signaling in population 2. (**C**) Expression of select TGF-β pathway members and associated genes in population 1-4 cells as detected in the RNA-seq analysis (n=3)(**D**) Expression of mesenchymal GBM subtype signature genes and proneual GBM subtype signature genes as detected in the RNA-seq of surface marker identified cell populations. (**E**) UMAP feature plot of CD51, CD63, and CD71 expressing cells illustrating the distribution of cells expressing each surface maker. (**F**) Distribution of CD51+CD63+CD71+ population 1 cells and CD51+CD63-CD71-population 2 cells in the UMAP plot illustrating enrichment in of triple positive cells in cluster Ast.3. (**G, H**) Heatmap of population 1 enriched genes (**G**) and population 2 enriched genes (**H**) among the 10 cell clusters identified in the scRNA-seq. One-way ANOVA *p<0.05, **p<0.01, ***p<0.001 followed by Tukey’s post-hoc test.

To investigate the molecular function and signaling pathways unique to each population, we performed gene ontology and pathway analysis of upregulated genes in populations 1 and 2 using the aggregated gene list enrichment analysis database Enrichr (Chen et al., 2013). Pathway analysis for population 1 revealed categories associated mostly with TGF-β regulation of extracellular matrix, which is consistent with the outcome of pathway analysis for cluster Ast.3 from the scRNA-seq. Notch signaling was the primary enriched pathway detected in population 2, which functionally aligns with the enrichments in neuron development observed in gene ontology analysis for cluster Neu.1 from the scRNA-seq (Fig 5B). TGF-β signaling induces proliferation of GBM and promotes migration (Pauklin and Vallier, 2013; Wang et al., 2018), and many TGF-β associated genes are upregulated in mesenchymal GBM tumors(Wang et al., 2016; Wang et al., 2018). Expression levels of TGF-β pathway members, including SMAD 2/3 as well as TGF-β associated genes such as TGFBR2, RUNX1, and TAGLN are highly expressed in population 1 compared to the other three populations in our bulk RNAseq data (Fig 5C). We also analyzed proneural and mesenchymal signature genes(Phillips et al., 2006; Verhaak et al., 2010; Behnan et al., 2019), and found that mesenchymal genes such as SERPINE1 and TIMP1 were enriched in population 1, whereas proneural genes such as DLL3 and STMN4 were enriched in population 2 (Fig 5D). These findings suggest that cellular pathways involved during normal astrocyte development may be recapitulated by cells in these tumors.

### Population 1 from cell surface marker analysis corresponds to cluster Ast.3 in the scRNA-seq

To determine whether cell populations 1 and 2 identified by the expression of CD51, CD63, and CD71 correlate with specific cellular clusters identified in the scRNA-seq analysis, we first examined the expression of the three surface markers in the scRNA-seq dataset. Feature plots revealed CD51, CD63, CD71 transcripts are enriched in astrocytic clusters Ast.1-3 (Fig. 5E), demonstrated both by individual gene expression and by the identification of cells that express all three surface markers (Fig. 5F). Quantification of triple-positive cells in clusters Ast.1-3 revealed that population 1 cells make up 31.52% of total cells in cluster Ast.3 but appear less frequently in Ast.1 (9.1%) and Ast.2 (4.74%) clusters. Even fewer triple-positive cells are detected in the neuronal clusters (Neu.1=1.56%; Neu.2=0.34%; Neu.3=0.22% Fig. 5F), in which surface marker expression is generally lower compared to that detected in the astrocytic clusters (Fig. 5E). In contrast, cells that express CD51 but low CD63 and CD71, the equivalent of population 2 cells, are detected more frequently in the neuronal clusters (Neu.1=1.88%; Neu.2=1.25%; Neu.3=2.33%) than the astrocytic clusters (Ast.1= 0.2%; Ast.2=0.87%, Ast.3=0.91%, Fig. 5F), although not specifically enriched in cluster Neu.1. These findings are consistent with what we expect from the results of bulk RNA-seq characterization of population 1 and 2 cells.

We next examined whether DEG identified in populations 1 and 2 are differentially expressed among the 10 scRNA-seq classified clusters. The top 50 DEG in populations 1 was found to be highly expressed in cluster Ast.3 (Fig 5G), whereas population 2-associated DEG are enriched in neuronal clusters with high expression detected in cluster Neu.1 (Fig 5H). Comparison of DEG that are shared between population 1 and cluster Ast.3 cells confirmed TGF-β signaling regulation of extracellular matrix, as well as focal adhesion and smooth muscle contraction, as the top three shared pathways in both experiments. On the other hand, comparison of DEG in population 2 and Neu.1 demonstrated no clear overlapping pathways, suggesting more divergent functions despite transcript level similarities. Taken together, these analyses confirmed the co-expression of CD51, CD63, and CD71 identifies an astrocytic progenitor population with mesenchymal characteristics.

### Heterogeneity of hPSC-derived astrocyte lineage progenitor populations

The transcriptomic differences between cells in populations 1 and 2 and their correlation with distinct glioma and developmental progenitor subtypes led us to ask whether population 1 and population 2 exhibit unique cellular properties such as morphology and proliferation. We FACS isolated population 1 and 2 NPCs at the LBF stage and monitored morphological changes under brightfield microscope for 1, 3, and 7 days after initial sorting (Fig. 6A). After 1 day, cells from population 1 show larger cell bodies with multiple processes, resembling the stereotypical astrocytic morphology (Fig. 6B, arrows). In comparison, the majority of cells from population 2 show unipolar and bipolar morphology, resembling radial glia progenitors (Fig. 6B, arrowheads). After 3 days or 7 days in culture, the cells in population 1 remained multipolar in morphology (Fig 6B, arrows), whereas some cells in population 2 changed from small, bipolar cells to multipolar cells resembling astrocytes (Fig. 6B, arrows). Cultures from both cell populations showed a small increase in cell number at day 3 and 7, indicating ongoing proliferation of the cultured cells (Fig. 6B).

**Figure 6.**
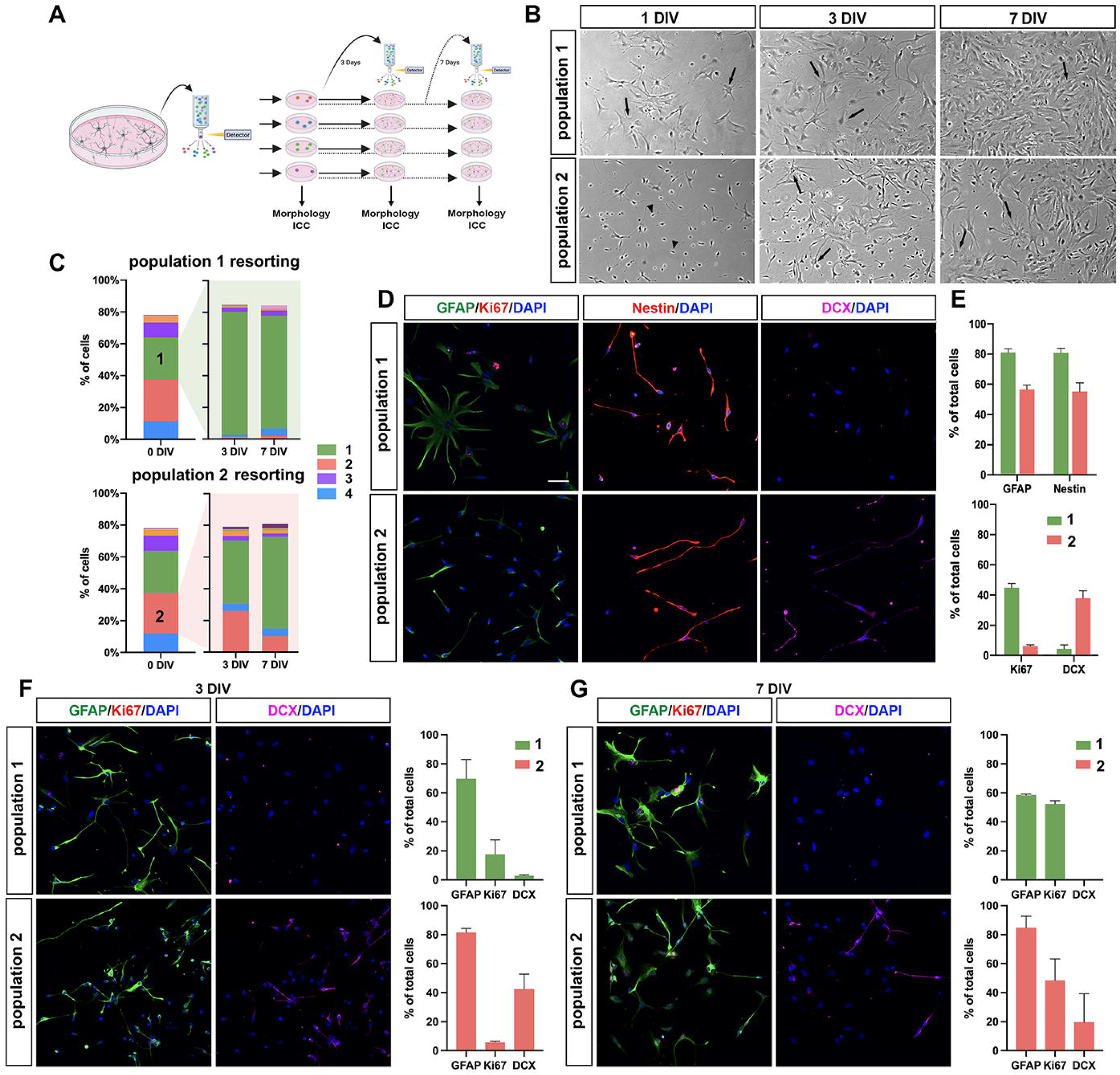
Characterization of GBM subtype-associated astrocytic progenitor populations reveal distinct morphological and proliferative properties. (**A**) Schematic of sequential FACS experimental setup and subsequent assays performed on the cell populations isolated based on CD51, CD63, and CD71. (**B**) Phase-contrast images of populations 1 and 2 cells at 1, 3, and 7 days *in vitro* (DIV) after sorting. (**C**) Quantification of the percentage of cell in populations 1 to 4 after sorted populations 1 and 2 were subsequently FACS isolated on day 3 and 7 (*n*=3). (**D, E**) Immunostaining representative images (**D**) and quantifications (**E**) of Nestin, GFAP, Ki67 and DCX-expressing cells in population 1 and population 2 at 1 DIV after FACS. (**F**) Immunostaining representative images and quantifications of cells expressing lineage markers GFAP, Ki67, and DCX after re-sorting populations 1 and 2 cells by FACS after 3 DIV and 7 DIV. *n* represents the number of independent differentiation experiments from one ESC line and two iPSC lines. Data are represented as means ± s.e.m. Scale bar, 50µm.

Next we used a combination of progenitor (Nestin), astrocyte (GFAP), and proliferation (Ki67) markers to verify the identity of cells in each population. The majority of the cells from population 1 and 2 expressed Nestin at day 1, indicating a progenitor phenotype (Fig. 6D, 6E). Quantification of GFAP+ cells in both populations 1 and 2 at day 1 revealed higher number of cells expressing GFAP in population 1, as well as striking morphological differences between GFAP+ cells found in each population (Fig. 6D, 6E). In addition, higher numbers of Ki67^+^ cells were detected in population 1 compared to population 2 (Fig. 6D, 6E), suggesting the presence of more cells actively in cell cycle in population 1. Consistent with our RNA sequencing findings, day 1 cultures from population 2 but not population 1 contained DCX expressing cells (Fig. 6D, 6E), indicating that population 2 had cells committed to the neuronal lineage.

To investigate whether combinatorial expression of the surface markers identifies specified or evolving cell types along the astrocytic lineage, we examined multipotency/interconversion capabilities of each population by re-sorting the FACS-purified populations over a period of 3 and 7 days (Fig. 6A, 6C). We found that cells in population 1 maintain their original surface marker profile as well as GFAP expression after 3 and 7 days (Fig. 6C), suggesting that cells in population 1 appear to be astrocytic lineage restricted despite a higher rate of proliferation as indicated by Ki67 quantification (Fig 6F, 6G). However, significant numbers of FACS purified population 2 cells acquire population 1 surface marker profile, express GFAP, and exhibit multipolar morphology at day 3 and 7. However, resorted population 2 cells maintain their original surface marker profile and express DCX at day 3 (Fig 6F) but appear to convert to population 1 by day 7 with increased Ki67 and reduced DCX expression (Fig. 6G). Taken together, our data suggest that population 2 contains a small population of bipotent progenitors that could commit to either neuronal or astrocytic lineages, whereas population 1 contains highly proliferative progenitors that are restricted to the astrocyte lineage.

## Discussion

Transcriptomic profiling of primary cells from adult and fetal human brains as well as cultured gliogenic neural progenitors have identified several molecularly distinct cell populations in the astroglial lineage (Zhang et al., 2016; Sloan et al., 2017; Weng et al., 2019; Barbar et al., 2020), but their relationships to the progenitor populations found in different GBM subtypes are not well understood. Couturier et al (2020) first compared progenitors in GBM tumors with primary neural progenitors from fetal brains at the single cell level, and identified cell populations in both sources that share gene expression profiles with TCGA GBM subtypes. In this study, we developed a hPSC differentiation protocol that produces the several progenitor subtypes during gliogenesis and identified cell populations that share transcriptomic similarities with tumor cells from the mesenchymal or proneural GBM subtypes. In addition, combinations of CD51, CD63, and CD71 were found to enrich for progenitors with mesenchymal or proneural properties. Taken together, these findings suggest hPSC-derived gliogenic progenitors generated under these experimental conditions may serve as a novel tool to investigate the developmental trajectory of GSC in specific GBM tumors.

Analysis of cell fate markers at two timepoints during our differentiation protocol revealed a molecular snapshot of the cells present during astrocyte development. At 160 DIV the cultures were heterogeneous with different populations of NPCs, astrocyte precursors, astrocytes, and a few neuronal progenitors. However, by 220 DIV the cultures were less heterogeneous with majority of cells expressing mature astrocyte markers. The gene expression profiles of the glial progenitors and astrocytes generated by our protocol are similar to those in HepaCAM-purified primary astrocyte lineage cells in adult and fetal human brains (Zhang et al., 2016). Further, progenitor populations identified at 160 DIV also correspond to distinct progenitor subtypes identified by scRNA-seq in the human fetal brain (Barbar et al., 2020), lending further confirmation that our differentiation protocol recapitulates the intricacies of human glial lineage commitment detected *in vivo*.

Combinations of the cell surface markers CD51, CD63, and CD71 were previously shown to label five subpopulations of ALDH1L1-expressing astrocytes from adult mice that correlated with GBM subtypes (John Lin et al., 2017). CD51, CD71, and CD63 are not known markers of human astrocyte precursors or astrocytes, but expression of all three proteins is increased in astrocytomas and glioblastomas (Rorive et al., 2010; Verhaak et al., 2010; Roth et al., 2013). We found that the transcriptome of human CD51+CD63+CD71+ cells (population 1) closely resemble that of the mesenchymal subtype of GBM, and that CD51+CD63-CD71-cells (population 2) resemble the proneural GBM subtype. By contrast, the mouse study found no cells that correlated with the proneural subtype, and that CD51+CD63-CD71+ cells (corresponding to population 4 in our study) were best aligned with the mesenchymal subtype, highlighting the marked differences between mouse and human glial lineages. There are also methodological differences that may explain some of the differences between this human study and the mouse study. The mouse study enriched for mature astrocytes by isolating Aldh1L1-GFP expressing cells from adult mouse brain whereas this study utilized developmental timing to enrich for glial progenitors without cell type-specific marker enrichment. The absence of a proneural subtype aligned population in the mouse study likely reflects the restriction of the analysis to only Aldh1L1-expressing cells.

Pathway analyses of the DEG in populations 1 and 2 also revealed differences in signaling pathways that correlate with findings in GBM subtypes. Activation of TGF-β signaling is detected in the mesenchymal subtype concurrent with a decrease of Notch signaling; conversely activation of Notch but not TGF-β signaling is seen in the proneural subtype (Saito et al., 2019). This matches what we found in populations 1 and 2 respectively. Similar enrichment in genes associated with TGF-β signaling was observed with cells in cluster Ast.3 identified in the scRNA-seq, which also expresses the highest levels of mesenchymal GBM and population 1 markers. It has been previously reported that the transcriptomic profile of mesenchymal subtype GBM has a resemblance to cultured astrocytes (Verhaak et al., 2010; Zong et al., 2012). ScRNAseq analyses of human fetal brain and tumor cells also demonstrated that cell clusters with mesenchymal properties share transcriptomic similarities with astrocytic clusters based on their proximity in UMAP dimensional reduction plots (Couturier et al., 2020). This similarity was also observed in our study, where mesenchymal Ast.3 cluster is adjacent to astrocytic Ast.1 cluster in the UMAP plot.

The lineage trajectory analysis also highlighted the divergent states of the cells expressing mesenchymal and proneural GBM markers, placing the two cell populations at distinct points along the glial differentiation trajectory. The localization of cells with a mesenchymal GBM signature at the distal end of the astrocytic lineage enriched path III as well as the left edge of the progenitor enriched path I suggests a population with both progenitor and mature astrocyte transcriptomic profiles. By contrast, localization of cells with the proneural GBM signature primarily in medial cell clusters of paths I and II suggests a population of intermediate progenitors with neuronal differentiation potential. Our lineage trajectory analysis is consistent with studies of human glioma GSC that revealed two distinct groups, a stem-like subtype that resembles a proneural signature and a precursor-like subtype that expresses mesenchymal signature (Cusulin et al., 2015). This correlates well with our finding that the proneural subtype-associated population has a more progenitor cell phenotype and that the mesenchymal subtype-associated population is more differentiated along the astroglial lineage trajectory. Importantly, expression of cell cycle markers such as MKI67 in the trajectories of both the mesenchymal and proneural-associated populations indicate that both populations possess proliferative potential. Ki67 immunostaining in sequentially FACS isolated populations 1 and 2 also suggested differential rates of expansion over time between the two populations, highlighting the cellular differences in these molecularly distinct progenitor subtypes. Despite the transcriptomic similarities between the hPSC-generated progenitors with mesenchymal or proneural properties with cell detected in respective GBM subtypes, our study is limited in demonstrating whether the hPSC-generated GBM-like progenitors are tumorigenic. Future xenograft studies will elucidate whether glioma-associated mutations in these two distinct cell populations lead to differences in tumor forming capacity in mouse models of GBM.

In summary, our study demonstrated that hPSC-derived astrocytic progenitors retain similar molecular heterogeneity as that observed in human fetal brain and GBM tumor cells. The ability to generate and isolate astrocyte progenitors with either mesenchymal or proneural-like properties from hPSC will facilitate our ability to decipher the differences between normal and malignant cells, and potentially devise new therapeutic approaches for the treatment of GBM.

---

**Extended data figure 1-1. Transcriptomic, molecular, and functional analyses revealed enrichment of mature astrocytes in long term culture.**

**(A)** Experimental timeline of astrocyte differentiation protocol including AMM time point. AMM = astrocyte maturation media. (**B**) Immunostaining of GFAP, S100b, and Tuj1 in astrocytes transitioned to AMM. (**C**) Quantification of Tuj1+, GFAP+ and S100b+ cells to total number of DAPI+ nuclei (*n*=3). (**D**) Immunostaining of mature astrocyte markers ACSBG1, GJA1 (Cx43), and SPARCL1 in astrocytes transitioned to AMM. (**E**) Quantification of ACSBG1+, GJA1+, and SPARCL1+ cells to total number of DAPI+ nuclei (*n*=3). (**F**) Expression analysis by quantitative PCR (qPCR) of astrocyte specific genes of cells at the NPC, LBF and AMM time points. Expression levels are presented as fold change relative to the NPC group (one-way ANOVA). DNA sequence of primers used for the qPCR analyses are listed in extended data table 1-1. (**G**) Glutamate uptake assay on NPCs and astrocytes from AMM stage. Bar graphs show concentration of glutamate taken up by hPSC-derived NPCs (n=3) and AMM astrocytes (n=5) after incubation with 50 mM glutamate for 1 h. (**H**) Astrocytes at the AMM stage respond to ATP. Representative traces of calcium transients from five astrocytes loaded with Ca^2+^ indicator Fura-2/AM following 100uM ATP application. Percent of total astrocytes exhibiting calcium response (*n*=5). (**I**) t-distributed stochastic neighbor embedding (tSNE) plots of single-cell RNAseq data from hPSC-derived, LBF-induced cells at 225 days in AMM stage (*n*=654). Four cell clusters were identified with cells with mature astrocyte profile (green) as the predominant cluster. (**J**) tSNE feature plots of astrocyte precursor genes (SOX9, NF1A, HOPX), astrocyte genes (GFAP, S100β, and HtrA1), cell-cycle genes (MKI67), and neuronal genes (DCX and SYN1), supporting the presence of four cell clusters including astrocytes, neuron, non-cycling bipotential intermediate precursors (IPCs), and cycling neural progenitors (NPCs). (K) Heatmap of top differentially expressed genes across the 4 identified cell clusters. Data in panels C, F, G, H are represented as means ± s.e.m. * p < 0.05; Scale bar = 50µm.

**Extended data table 1-1. DNA sequence for primers used in quantitative PCR validation of cell fate markers at the AMM stage.**

**Extended data figure 2-1. Heatmap of GBM subtype enriched genes identified cell clusters with similar expression profiles.** (**A**) Heatmap representation of relative expression levels of mature and fetal astrocyte enriched genes as identified in Zhang et al., 2016 (see complete list in Extended data table 2-2). (**B**) Heatmap of classical GBM subtype markers from Verhaak et al., 2010 in the 10 clusters identified in the scRNA-seq of LBF-induced cells. Marker expression is most frequently detected in cluster Ast.1 (A1) but also detected in clusters Ast.3 (A3) and Neu.1 (N1). (**B**) Heatmap representation of transcript levels of mesenchymal, proneural and classical GBM subtype markers identified in Wang et al., 2017. Similar enrichments of mesenchymal markers in cluster A3, proneural markers in cluster N1, and classical markers in cluster A1 confirmed the results obtained using subtype markers from the Verhaak study.

**Extended data table 2-1. Cell clusters and cluster-specific DEGs from scRNA-seq analysis of LBF stage progenitors.**

**Extended data table 2-2. Gene lists for fetal vs mature astrocytes and the different GBM subtypes used for heatmap analyses.**

**Extended data table 4-1. Cell population-specific DEGs in populations 1 to 4 in pairwise and 1 vs. 3 comparisons.**

