## Extended Data Table 1-1 for "Identification of Human Pluripotent Stem Cell Derived Astrocytic Progenitors that Correlate with Glioblastoma Subtypes"

**Extended data table 1-1. Quantitative PCR Primers Used for the Study**

| Genes | Type | Sequence (5’-3') |
| --- | --- | --- |
| HtrA1 | Forward | CAAAGTTGAGCTGAAGAACGGT |
|  | Reverse | TGGTCAATTTTGATGAGTGCGAT |
| S100b | Forward | TGGCCCTCATCGACGTTTTC |
|  | Reverse | ATGTTCAAAGAACTCGTGGCA |
| GAPDH | Forward | ACACCATGGGGAAGGTGAAG |
|  | Reverse | GTGACCAGGCGCCCAATA |
| SPARCL1 | Forward | GCACCTGACAACACTGCAATC |
|  | Reverse | TTTCAGCCTTATGGTGGGAATC |
| GJA1 | Forward | AGTACCAAACAGCAGCGGAG |
|  | Reverse | CTCCAGTCACCCATGTTGCC |
| GFAP | Forward | TCCTGGAACAGCAAAACAAG |
|  | Reverse | CAGCCTCAGGTTGGTTTCAT |
