## Supplementary figures and images for "Identification of Human Pluripotent Stem Cell Derived Astrocytic Progenitors that Correlate with Glioblastoma Subtypes"

### Extended Figure 1-1

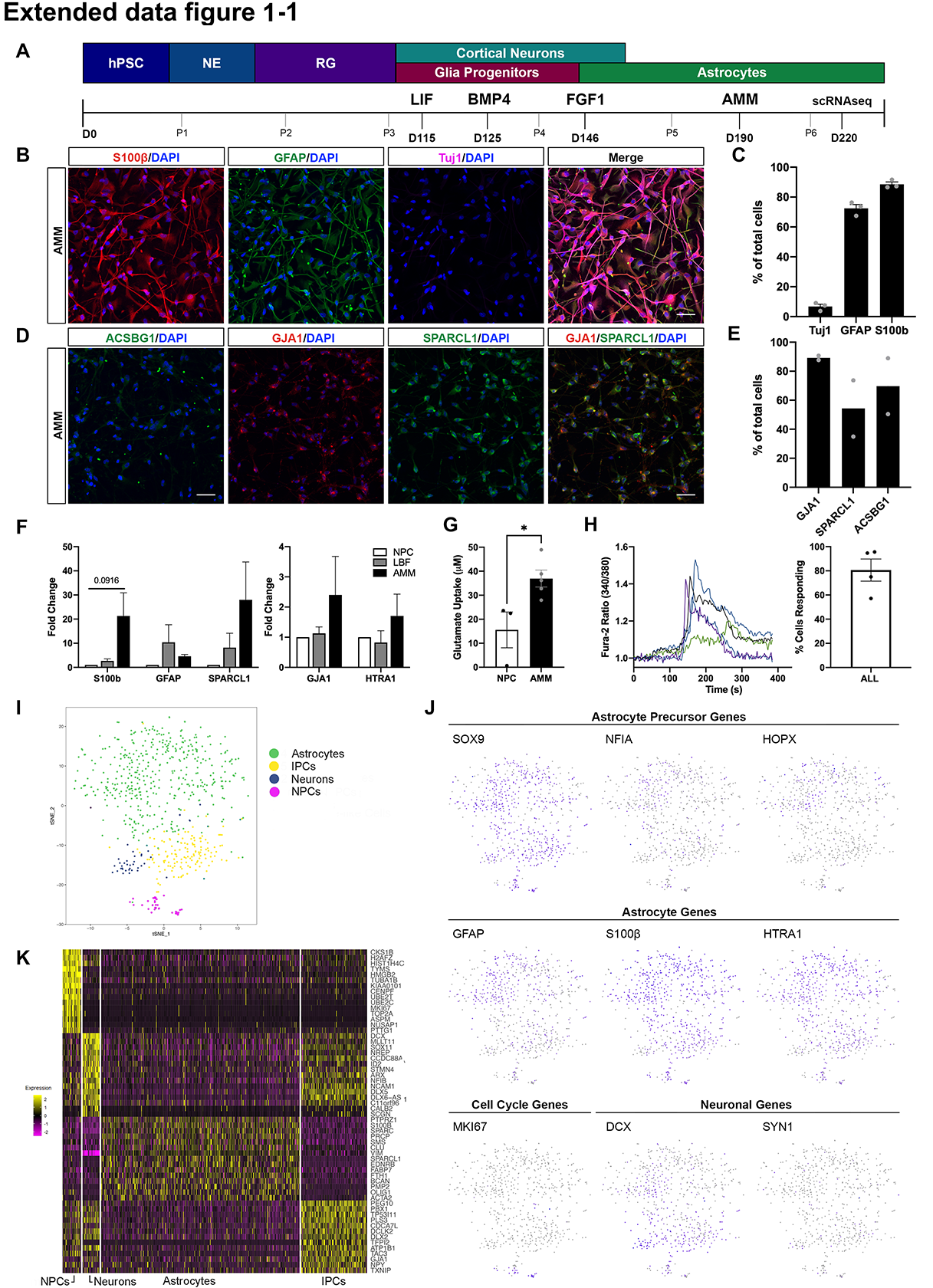

### Extended Figure 2-1

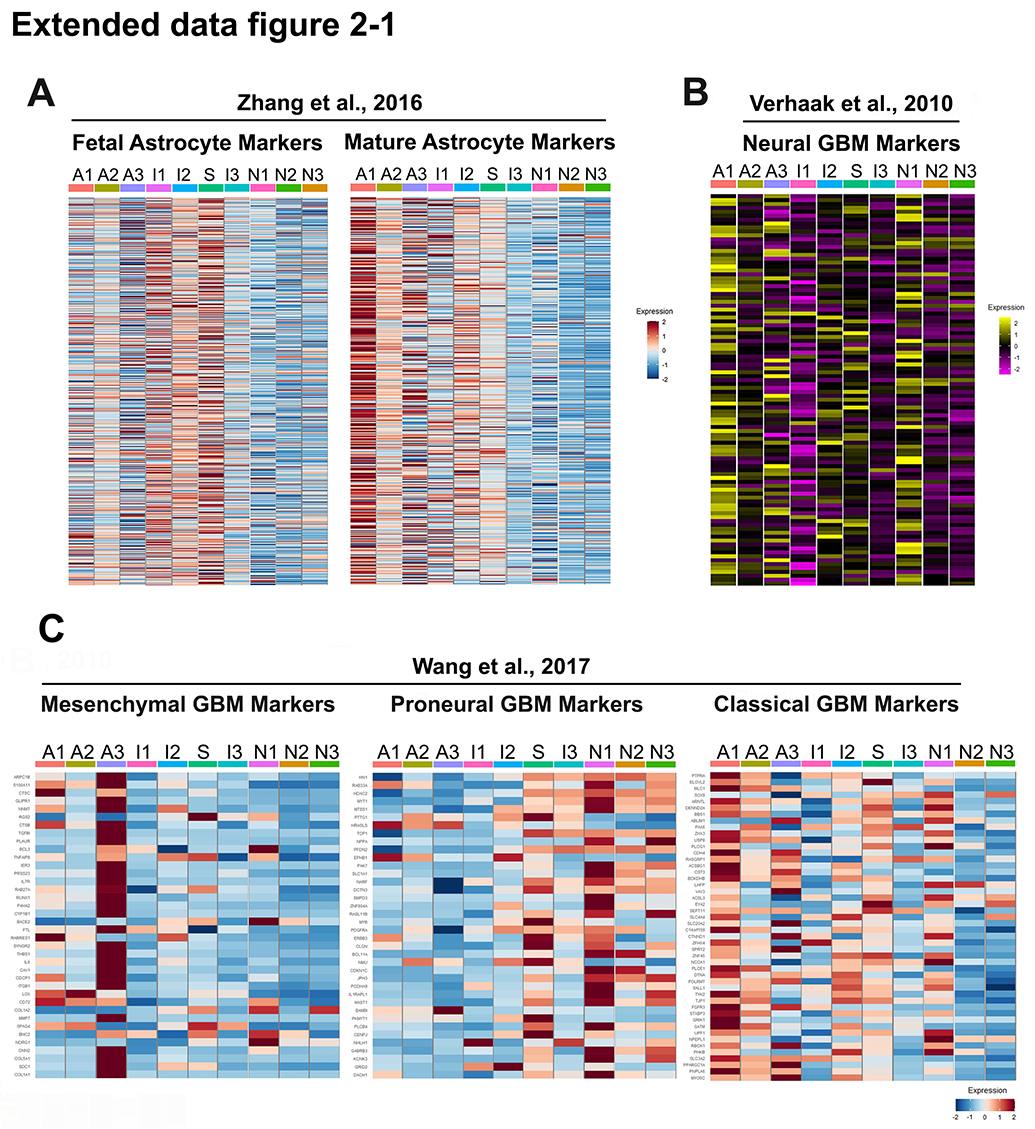
